# Vampire bats target their social grooming to hard-to-reach body parts

**DOI:** 10.64898/2026.02.25.707567

**Authors:** Chuanliang Chen, Tobias I. Nguyen, Michael Meyer, Evan Hashem, Gerald G. Carter

**Affiliations:** Department of Evolution, Ecology and Organismal Biology, The Ohio State University, Columbus, Ohio, United States of America; Department of Ecology and Evolutionary Biology, Princeton University, Princeton, New Jersey, United States of America; Howard Hughes Medical Institute, Chevy Chase, Maryland, United States of America; Smithsonian Tropical Research Institute, Balboa, Ancón, Panamá

## Abstract

Many group-living mammals and birds groom the fur (or preen the feathers) of their close associates to build and maintain affiliative relationships. Several lines of evidence suggest that social grooming in female common vampire bats (*Desmodus rotundus*) is a low-cost form of helping, but it has yet to be shown that vampire bats preferentially groom others in locations that are difficult to self-groom, as has been found in multiple primate species. In this short note, we show that vampire bat social grooming (n = 1586 events) occurs most often on parts of the recipient’s body where self-grooming (n = 1515 events) is least likely, and often in locations where the recipient could not lick itself, like the back of the head. The finding that vampire bats preferentially groom each other in hard-to-reach locations provides further support for the hypothesis that social grooming is both a social signal and a low-cost form of helping.

## Introduction

In many group-living birds and mammals, and especially primates, individuals groom the fur (or preen the feathers) of group members or preferred close associates [1–4]. Like other forms of “social touch” [5], this social grooming (or allogrooming) signals affiliative intentions, but it also provides cleaning services that supplement self-grooming [6–10] by cleaning wounds [11], removing harmful ectoparasites [9,12–16], and smoothing out fur and feathers for shedding water and holding heat [8,17]. In primates, individuals often invest substantial time and energy grooming others, suggesting that social grooming has a “dual function” [9] as both a low-cost cooperative investment and a signal of a groomer’s willingness to cooperate in other ways that provide access to food or social support [18–23].

Female common vampire bats (*Desmodus rotundus*) spend about 5% of their awake time licking the fur or wings of other bats —a greater frequency of social grooming than observed in any other bat species [24]. Previously unfamiliar and unrelated vampire bats use reciprocal grooming to ‘test the waters’ while forming new reciprocal food-sharing relationships [19], in which partners regurgitate ingested food (blood) to feed each other when one has failed to feed [25,26]. Unlike food sharing in times of need, social grooming can be reciprocated immediately, making it a low-cost and low-risk way to both build up trust and immediately assess a cooperative relationship [19].

Social grooming in vampire bats appears to benefit recipients and be responsive to cues of need, such as experimentally wetted and disturbed fur [27]. If social grooming is indeed a low-cost form of helping, then individuals should preferentially groom each other in locations that are difficult to self-groom, a pattern that has been reported in multiple group-living primates and other social species (reviewed in “Discussion” below), but not yet in vampire bats. In this short note, we show that vampire bats do preferentially groom each other in locations that are difficult to self-groom.

## Materials and Methods

To compare locations on the body for both self-grooming and social grooming events, we first sampled 1586 previously recorded social grooming events from 61 sampled hours across 17 days of recorded surveillance videos of interactions among common vampire bats (*D. rotundus*) housed in a flight cage at the Smithsonian Tropical Research Institute in Gamboa, Panama (26 adult females, 2 adult males, and 1 juvenile male; for details of the colony, see Razik et al. [28]). To avoid ambiguity, we only recorded social grooming events that were at least 5 seconds in duration. For each social grooming event, we labeled the location on the body that was groomed for the most time during the event as being one of 11 locations: face, head, upper back, lower back, chest, belly, foot, arm, thumb, inside wing, and outside wing (Fig. 1). The leg was initially included as a location but received only one self-grooming event and no social grooming events, so we excluded it from the analysis.

**Figure 1.**
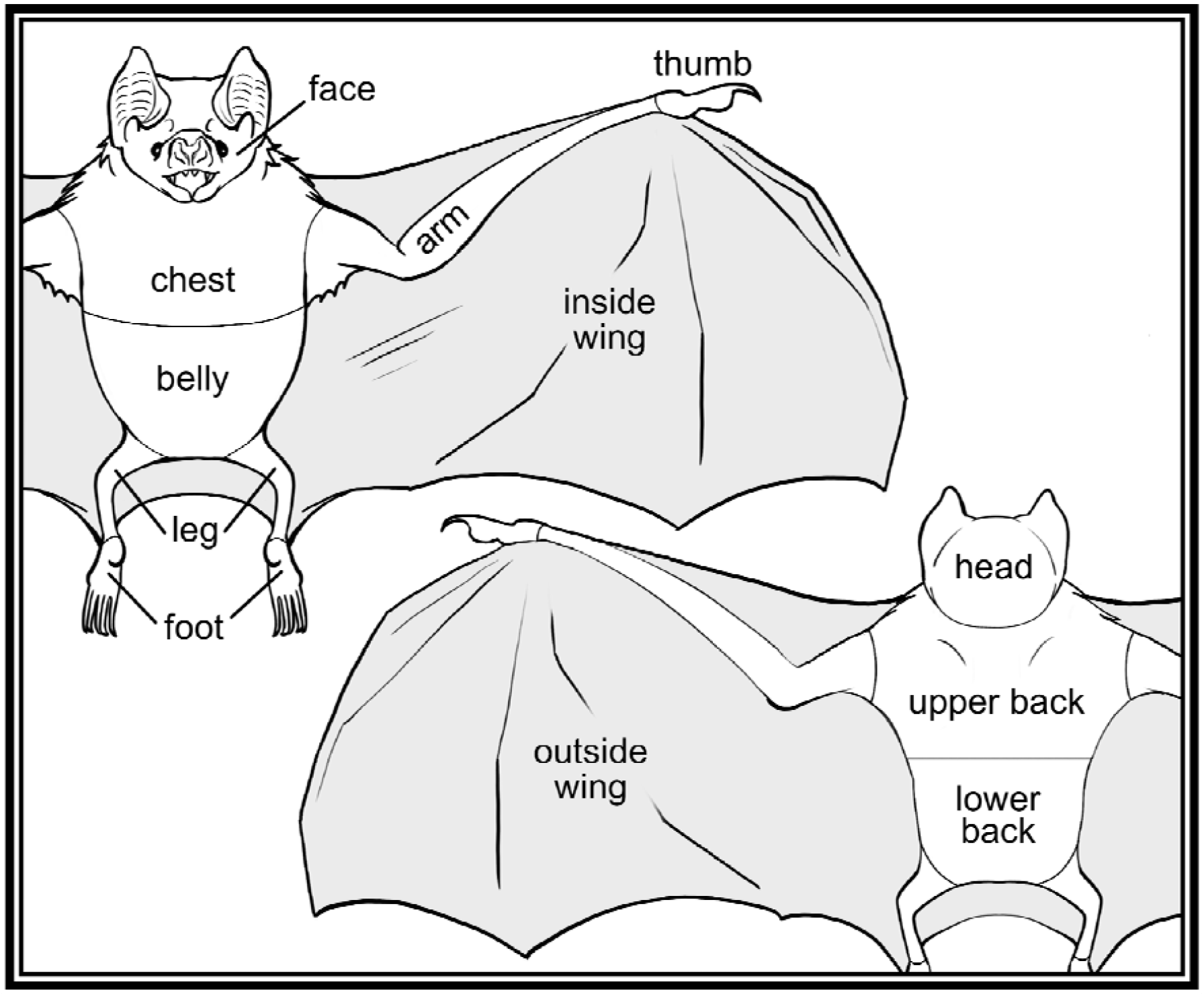
Locations for self-grooming and social grooming events.

To ensure social grooming and self-grooming events were sampled at the same times of day, we attempted to find a paired observation of self-grooming for each social grooming event (i.e. the next self-grooming event to be observed by any bat on camera), and we labeled the self-groomed body location, leading to 1515 self-grooming events. To assess which body locations were less accessible to be self-licked, we also recorded whether 1128 of the self-grooming events were performed by either licking or scratching.

We used a permutation test to determine whether the distributions of self-grooming and social grooming were more convergent or divergent across the body than expected by chance. To do this, we calculated the absolute difference between the proportion of total self-grooming that occurred on a given body part and the proportion of total social grooming that occurred on that same body part, then averaged these differences across all 11 body parts. We then compared the observed value to the distribution of 5000 expected divergence values generated by repeatedly shuffling the locations across all grooming events, which preserves the frequency of locations but removes any association with type of grooming. We calculated the p-value as the percentage of the expected values that were more extreme than the observed statistic.

## Ethics statement

This work was approved by the Smithsonian Tropical Research Institute Animal Care and Use Committee (no. 2015-0501-2022) and the Panamanian Ministry of the Environment (no. SEX/A-67-2019).

## Results

Social grooming occurred most often on parts of the recipient’s body that were the least likely to be self-groomed (Fig. 2, observed divergence = 12.5%,95% quantiles of expected values = 0.4% to 1.2 %, p < 0.0002, Fig. 3). The bats used almost exclusively licking to groom their inside wing, foot, thumb, and arm, and they almost exclusively used scratching with their foot to clean their belly, lower back, chest, upper back, head, and face (Fig. 4). A bat’s face, back of head, back, and chest could only be reached by scratching, and these were also the locations most targeted by allogroomers (Fig. 2).

**Figure 2.**
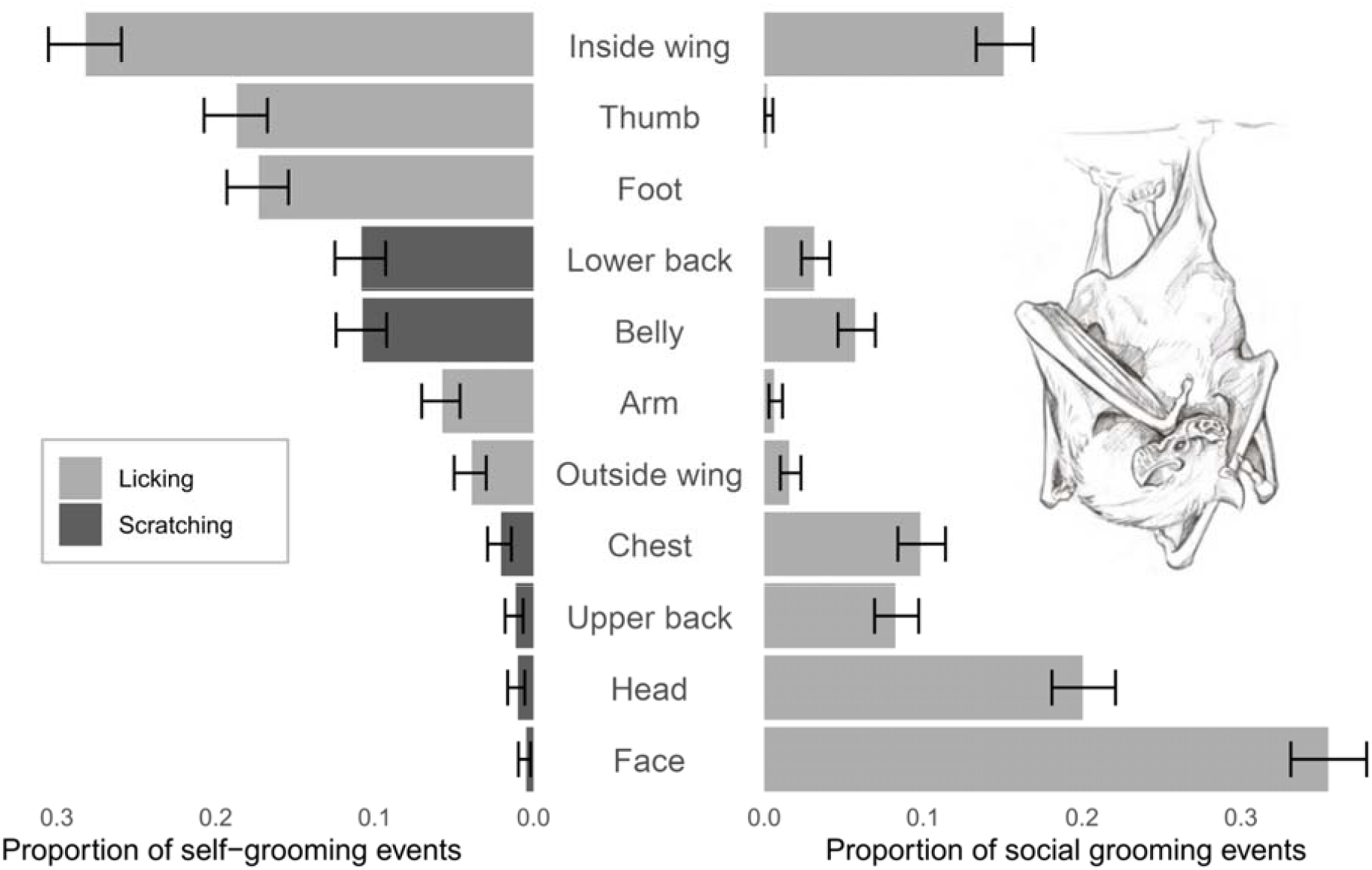
Self-grooming and social grooming are targeted to different body locations. Locations on the body (y-axis) are listed in order of self-grooming frequency. Error bars show 95% confidence intervals from a binomial test. Bat illustration by Javier Lazaro.

**Figure 3.**
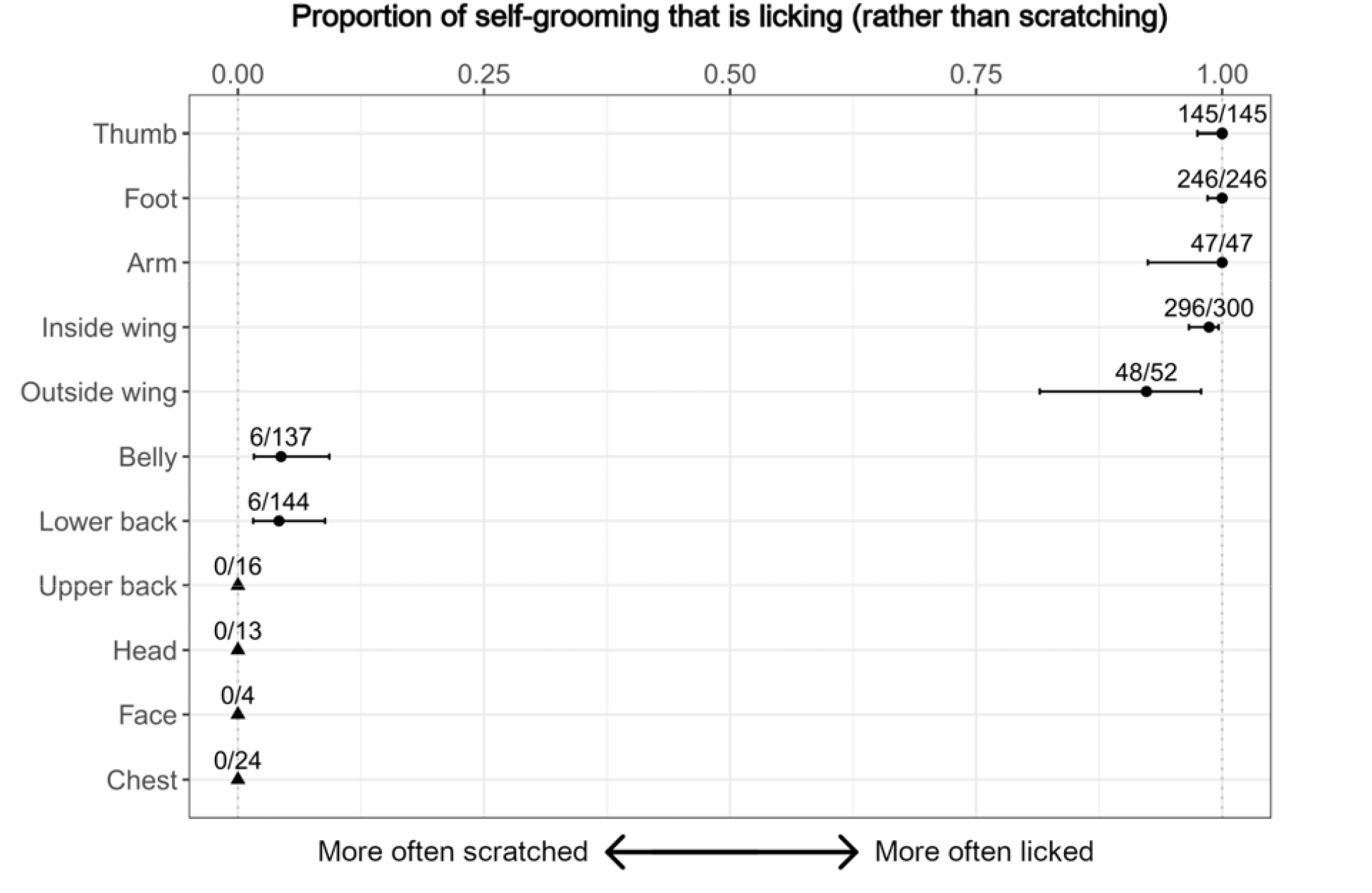
Vampire bats groomed body parts by either licking or scratching but not both. Points are proportions of a sample of self-grooming events that were licking for each body location (0 is 100% scratching and 1 is 100% licking). Numbers show counts of licking out of total cases. Error bars are 95% confidence intervals from a binomial test, except for locations where self-licking is impossible (triangles).

## Discussion

Female vampire bats preferentially groom each other on body locations that are the least self-groomed and that appear difficult or impossible for the recipients to lick by themselves, such as the back of the head, face, and upper back. This finding corroborates other observations that social grooming in vampire bats is a form of low-cost helping that is sensitive to cues of need. For example, bats receive more social grooming when they have experimentally wetted and disturbed fur and immediately after they have groomed themselves [27]. Anecdotally, it is clear that social grooming is also targeted to novel foreign objects like metal bands on the forearm.

One might argue that the distribution of social grooming we observed resulted from the fact that many body parts that are hard to self-groom also happen to be most accessible to the groomer when hanging face-to-face. However, the amount of social grooming on ventral body parts (chest, belly) was only slightly more than their respective dorsal counterparts (upper and lower back), even though the ventral side is more accessible to a face-to-face social groomer. Moreover, bats directed very little social grooming to the thumbs, which are highly accessible in that position. Vampire bats have large, specialized thumbs that they use to jump, land, walk, and run [29–32]. The most obvious reason the thumbs would not be socially groomed is that they receive frequent self-grooming.

The only exception to our finding that social grooming occurs where self-grooming is rare is the fact that the inner wing was frequently groomed during both self- and social grooming. This finding cannot be explained by the large surface area of the inner wing, because the outside wing membrane has the same surface area and received little self- or social grooming. Instead, the inner wing may be socially significant or require extra grooming for reasons we do not yet understand.

In our study, the most socially groomed body part was the face. In some primates, it has been suggested that the face receives frequent social grooming in part because of its role in communication and individual recognition [33–35]. It is possible that some small amount of grooming on the face was due to mouth licking, which serves as a means to share food, a signal of intent to share or receive food, and possibly for gathering information or sharing microbiota [36]. However, the bats in our study were not fasted and mouth licking among non-fasted bats is very infrequent [36].

Our finding that vampire bats preferentially groomed each other in locations that are difficult to self-groom has been reported in multiple other species, including chimpanzees [3,10], bonobos [37], 5 lemurs [3,13,38], 24 other primate species [3,6,13,17,33,39–47], bird species from over 35 families [2,9,48], impala [12,49], cattle [7], mule deer [50], the European badger [51], and honey bees [52,53]. Some of these studies also suggest that self-scratching is inferior to social grooming by licking [e.g. 12]. In rockhopper and macaroni penguins, social grooming mainly occurred between mated pairs; accordingly, unpaired individuals hosted more ticks and more frequently scratched their heads with their feet [54]. A study of four antelope species found that all species scratched with their hind legs, but this behavior was less frequent in the impala [12], a species which engages in regular grooming of the head and neck [49,55].

The degree of cooperation or conflict in a relationship might be reflected by the degree of coordination of movement to facilitate or prevent allogrooming. In many other species, the recipient exerts control over the site of social grooming by moving to expose different parts of its body [6,39,52,55–59]. In vampire bats, for example, we noted that the inside and outside wing could only be groomed when the recipient opens its wing. Across species, some body sites require more cleaning, due to longer hair [6,41], being more frequently soiled [3], wounds [11], or ectoparasites [15]. In vervet monkeys, the recipient presents new body parts possibly to prolong the grooming session [60], and in three species of macaque, the groomer apparently switches to grooming the back, tail, or upper leg to communicate the end of the social interaction [6,61]. In primates, grooming face-to-back rather than face-to-face is thought to prevent eye contact, protect the groomer from biting, and protect the recipient’s vulnerable ventral side [34,37,40,44,56,62]. For this reason, the back and tail may receive more frequent grooming when the risk of aggression is increased, such as when there is a difference in rank (e.g. rhesus macaques [56], long-tailed macaques [34], mandrills [62], bonobos [37], and the langur *Semnopithecus entellus* [40]), when the groomer is male [34,37], when the recipient is carrying an infant [56], or when agonistic behavior preceded the grooming bout [44,56]. Bonobos more frequently groomed face-to-face than chimpanzees, which are thought to be more aggressive [63], and the same pattern was found with bonnet macaques versus the more aggressive pigtail macaques [6]. Future studies in vampire bats should assess whether coordination and locations of social grooming differ across relationships that vary in strength or conflict.

## Supporting information

R script for data analysis

## Acknowledgements

We thank Grace Vance and Imran Razik for help with data collection. We thank Rachel Page for access to resources.

## Supporting information

### S1 File. Raw data

### S2 File. R script for data analysis

## Notes

### Competing Interest Statement

The authors have declared no competing interest.

### Summary of Updates

Introduction and discussion condensed; histogram of expected vs observed values removed.

### Funding

- U.S. National Science Foundation: IOS-2015928

